# Cancer-associated synonymous mutations reveal stress signal-dependent mRNA folding that selectively modulates protein function

**DOI:** 10.64898/2026.01.26.701754

**Authors:** Sivakumar Vadivel Gnanasundram, Lixiao Wang, Sa Chen, Ondrej Bonczek, Borivoj Vojtesek, Robin Fahraeus

**Affiliations:** Department of Medical Biosciences, Umea University, 90185 Umea, Sweden; Department of Biological Sciences & Engineering, Indian Institute of Technology Palakkad, Palakkad 678623, India; Inserm U1131, 27 Rue Juliette Dodu, 75010 Paris, France; RECAMO, Masaryk Memorial Cancer Institute, Zluty kopec 7, 65653 Brno, Czech Republic

**Keywords:** RNA structures, synonymous cancer mutation, DNA damage response, p53

## Abstract

Recent technical advances have facilitated studies on changes in mRNA structures in response to signaling pathways. However, if mRNA structures can affect the function of the encoded protein remains poorly understood. In-cell RNA structural probing (SHAPE-MaP) demonstrates how two cancer-associated synonymous mutations (CASMs) at proline codon 34 (c.102 C>A and c.102 C>G) prevent DNA damage-induced *TP53* mRNA folding, whereas the non-cancer-associated c.102C>U mutation does not. Transcript and chromatin immunoprecipitation (ChIP) analysis reveal that p53 expressed from CASM34 has reduced promoter binding and reduced induction of p53 downstream target genes *PUMA* and *14-3-3-σ*, but not *p21*^*CDKN1A*^. Transcriptome analysis reveals a *CASM34*-mediated global attenuation of DNA damage–responsive gene expression. Together, the results demonstrate that CASM34 interferes with signal-induced *p53* mRNA folding during DNA damage, leading to selective modulation of protein activity. More broadly, our findings highlight a general concept by which cancer-associated synonymous mutations target signal-induced mRNA structures that influence the encoded protein.

**Significance statement:** Recent studies have revealed that synonymous mutations can target RNA metabolism and are associated with numerous diseases; however, how these mutations affect the function of the encoded protein remain unclear. Using in-cell RNA structural probing, we show that two cancer-associated synonymous mutations in *TP53* disrupt DNA damage–induced folding of *p53* mRNA. These mutations selectively impair p53 promoter binding and transcriptional activation of specific downstream target genes. Our results demonstrate that disruption of signal-induced mRNA folding by synonymous mutations can modulate the activity of the encoded protein. More broadly, this work identifies signal-responsive mRNA structures as functional targets of cancer-associated synonymous mutations.

## Introduction

Genomic sequencing has increased the spectrum of cancer-associated mutations, but the assessment of mutation-linked biological effects remains a challenge. Cancer genetics primarily focuses on somatic missense and nonsense mutations that affect the encoded protein sequence (1, 2). For example, mutations that target the p53 tumour suppressor protein and its capacity to respond to various cellular stress-induced signaling pathways are seen in over half of human cancers (3). The majority of these affect its DNA binding activity and the expression of a plethora of downstream target genes. In response to DNA damage, these include cell cycle regulatory genes such as *p21*^*CDKN1A*^, *14-3-3-σ, and Gadd45* while factors like *NOXA, PUMA, Bak*, and Bax induce apoptosis if the damage is too severe (3–6).

Synonymous mutations (SMs) alter the DNA and RNA sequences without affecting the encoded protein. Despite compelling evidence supporting their association with multiple human diseases, including cancers, these are under-researched due to the dearth of the underlying molecular mechanisms on how they affect the function of the encoded protein (7–11). Cancer-associated SMs (CASMs) are found at higher frequencies in the *TP53* gene compared to other tumor suppressors. For example, the *TP53* CASM22 interferes with the formation of an MDM2-binding platform, impairs ATM-mediated phosphorylation of the nascent p53 at serine 15 and attenuates the induction of p53 expression during DNA damage (12–15). Moreover, the CASM203 mimics PERK kinase-induced RNA structures and promotes p53/47 isoform expression without activation of the unfolded protein response (16–18).

Some of the better-known examples of biological functions controlled by regulated RNA structures come from prokaryotic riboswitches (19) or viral RNAs (20). More recent advances in highly reproducible experimental techniques probing in-cell RNA structures are illustrating how mammalian mRNAs respond with specific structural changes in response to cell signaling pathways (15, 16). This is an emerging field with new techniques being developed that together with advanced computational models hold promise to integrate RNA structures with the overall cellular responses to intra- and extra-cellular changes in the cellular environment (21, 22).

Here we show that two *TP53* SMs in codon 34 identified in multiple cancers selectively affect the p53 DNA damage response (DDR). In-cell RNA structure probing demonstrates that the CASM34s prevent *p53* mRNA folding during DDR. Transcriptome-sequencing and genome-wide ChIP-seq show that CASM34s selectively affect p53 downstream target genes by reducing p53 promoter binding. Overall, our findings illustrate how stress signal-regulated *p53* mRNA structures selectively differentiate the activity of the encoded protein and how these structures are targeted by synonymous mutations.

## Materials and methods

### *Cell culture, transfection, and treatment*s

p53-null H1299 cells (non-small-cell lung carcinoma human cell line) were mostly used for experimental analysis. Cells were cultured in RPMI 1640 medium (31870074, Thermo Fisher Scientific) supplemented with 10% fetal bovine serum (A3160502, Thermo Fisher Scientific), 100 U.mL-1 penicillin and 100 mg. mL-1 streptomycin (15140122, Thermo Fisher Scientific) and 2 mM L-glutamine (25030081, Thermo Fisher Scientific) and maintained at 37°C in a humidified 5% CO2 incubator. Cell lines were routinely checked for mycoplasma contamination using MycoStrip™ - Mycoplasma Detection Kit (rep-mys-10, Invivogen). Plasmid DNA transfections were performed using GeneJuice reagent (70967, Sigma-Aldrich) following the manufacturer’s protocol. To induce DNA damage, cells were treated with 1 µM concentration of doxorubicin (doxo) (D1515, Sigma-Aldrich) prepared in Dimethyl sulfoxide (DMSO) (276855, Sigma-Aldrich) for 16 h, unless specified otherwise.

### Plasmid constructs

All constructs were generated using the pcDNA3 eukaryotic expression vector (Life Technologies, Carlsbad, CA, USA) unless stated otherwise. CASM34 mutations were inserted into the p53 coding sequences using site-directed mutagenesis and verified by Sanger sequencing.

### Western blotting

Cells were washed with ice-cold phosphate buffer saline and lysed in RIPA buffer (Thermo Fisher Scientific), supplemented with a complete protease inhibitor cocktail (Roche, Basel, Switzerland). Equal protein amounts were resolved in 10% Bis–Tris Plus Gels (Thermo Fisher Scientific), transferred onto the BioTrace NT pure nitrocellulose blotting membrane (PALL Corporation) and blocked with 5% non-fat dry milk in Tris-buffered saline pH 7.6 containing 0.1% Tween-20. Proteins were probed with corresponding antibodies (listed below) and detection was performed using WestDura (Thermo Fisher Scientific) with myECL Imager (Thermo Fisher Scientific). Antibodies: HRP-conjugated anti-p53 DO1 mouse mAbs (prepared in house); anti-actin mouse pAbs (AC-15, Sigma-Aldrich), Western blots represent n≥3 and the original uncropped blots are provided in the supplementary information.

### RNA isolation and qRT-PCR

The total RNA was purified from H1299 cells post-transfection using the RNeasy Mini Kit (74104, Qiagen) following the manufacturer’s protocol. RT was carried out using Superscript II Reverse Transcriptase (18064014, ThermoFisher Scientific) and oligo(dT) primers (18418012, ThermoFisher Scientific). RT-qPCR was performed on QuantStudio™ real-time PCR system (Applied Biosystems) using PowerUp™ SYBR™ Green Master Mix (A25741, ThermoFisher Scientific) with the corresponding p53 target primer sets (supplementary information).

### In-cell SHAPE-MaP

SHAPE-MaP was performed as described previously (16, 23, 24). Briefly, H1299 cells grown in 6 well plates were transiently transfected with indicated constructs and treated with the indicated conditions (DMSO/DOXO). 36 h post-transfection, cells were washed with PBS. The SHAPE reagent 1-Methyl-7-nitroisatoic anhydride (1M7) (Sigma-Aldrich) was added to a final concentration of 10 mM by adding 100 µL of 100 mM 1M7 to 900 µL of RPMI media and treated for ∼90 seconds at 37°C. The same volume of DMSO was added to the untreated samples. Cells were then washed with PBS and harvested. RNA purification was carried out using the RNeasy kit (74104, Qiagen), followed by DNAse I digestion for 30 minutes at 37°C. Reverse-transcription of purified RNA was carried out with the p53 specific primers(16), using the MaP buffer and Superscript II Reverse Transcriptase (18064071, Thermo Fisher Scientific). Synthesized cDNAs were then purified and amplified using Q5 DNA polymerase (NEB) using the indicated p53 primers. The PCR product was purified and quantified with a Qubit fluorometer and diluted to 0.2 ng/µL. Purified amplicons were then tagmented and a library was created using the Ilumina Nextera XT DNA library preparation kit. Products from library PCR were then purified with the Agencourt AMPure XP beads (A63880, Beckman Coulter Life Sciences) as described in(23). Library concentration was measured with a Qubit fluorometer and the size distribution was measured using an Agilent 2100 Bioanalyzer according to the manufacturer’s instructions. Libraries were then sequenced with the Paired-end 150 NovaSeq System (NovaSeq PE150, Novogene, UK). SHAPE reactivity profiles and comparisons were generated using the ShapeMapper 2 and deltaSHAPE scripts, using default settings as described in (23) and aligned to indicated p53 coding sequences (CDS). SHAPE-Map data shown are representative of at least two independent biological repeats. The variations across biological repeats of each experiment were examined by Spearman’s rank-order correlation coefficient (supplementary information), demonstrated excellent correlation across biological replicates.

#### Transcriptome sequencing

Total RNAs were isolated from H1299 cells expressing p53-WT/CASM34 using RNeasy Mini Kit (74104, Qiagen) following the manufacturer’s protocol. Quality of isolated RNAs were assessed by Agilent 2100 Bioanalyzer according to the manufacturer’s instructions. RNAs were then subjected to polyA purification, library preparation and massive parallel sequencing (Novogene, UK). RNA-Seq alignment, assembly, and quantification were carried out using STAR (2.7), RSEM (v1.2), and DESeq2 (v 1.44). Functional enrichment analysis of the differentially expressed genes (DEGs) were performed using the R package clusterProfiler 4.0 (25) and the web-based tool Enrichr (26).

### DNA-ChIP

p53-ChIP was performed using the Zymo-Spin ChIP kit (D5210, Zymo Research) following the manufacturer’s instructions. Briefly, harvested cells were washed with ice-cold PBS and resuspended in 1X PBS. Cell suspension was then cross-linked with 1% formaldehyde for 10 minutes at room temperature and stopped by addition of 125 mM glycine. Cross-linked cell pellets were subjected to nuclei preparation, chromatin shearing by sonication. Sheared chromatin was then subjected to IP overnight using DO-1 p53 antibody (2mg/ml) or IgG antibody control, 15 µl of ZymoMag Protein A beads were added each ChIP reaction and incubated for 2h at 4°C. The samples were then washed with low and high salt buffers and then eluted. Eluted DNA was de-cross-linked with high-salt buffer at 65°C for 1h and digested with proteinase K at 65°C for 2h. ChIP DNA was then purified using Zymo-Spin™ IC Column. qPCR was performed on purified ChIP DNA using the primers of indicated p53-downtream targets (supplementary information).

### ChIP-seq

p53 ChIP-seq was performed using a SimpleChIP® Plus Enzymatic Chromatin IP Kit (9005, Cell Signalling technology) following the manufacturer’s instructions. Briefly, cells were fixed and nuclei was purified as described in DNA-ChIP. DNA was then fragmented by treating with Micrococcal Nuclease for 20 min at 37°C and sonicated. Fragmented DNA was subjected to IP as described above using DO-1 p53 antibody (2mg/ml) and ChIP-Grade Protein G Magnetic Beads. Quality of the purified ChIP DNA was analyzed by Agilent 2100 Bioanalyzer according to the manufacturer’s instructions and proceeded for library preparation and deep sequencing using PE150 strategy on Ilumina platform (Novogene, Cambridge, UK). The paired end reads were trimmed to remove adaptors with Trim Galore (v 0.6) and aligned to the human genome (GRCh38) using Bowtie2 (v 2.3). BigWig coverage files were generated from duplicate-removed BAMs using deepTools bam coverage with RPKM normalization. MACS2 (v 2.1) was used to identify the peaks in the ChIP-seq data. HOMER was used to perform de novo and known motif enrichment on the MACS2-called peaks from the ChIP-seq data.

Statistical analysis. Statistical significance was analyzed by comparing data sets with corresponding reference points using two-tailed unpaired t test (*P < 0.05; **P < 0.01; ***P < 0.001; ns: not significant). Spearman’s rank-order correlation coefficient was used to assess the biological replicates of RNA SHAPE-MaP data sets. Statistical assessments were performed using GraphPad Prism software and Python.

## Results

### Two CASM34 mutants impose similar *p53* mRNA secondary structures

We shortlisted *Tp53* CASMs from p53 mutation (27) and SynMICdb (9) databases and assessed their potential effect on *p53* mRNA structures using in silico RNA structure analysis tools and cellular function(9, 28). From this we identified two mutants at codon 34 (proline) (CASM34 c.102 C>A and c.102 C>G) derived from malignant glioma and melanoma patients, respectively **(Figure 1A)**. We employed the high-throughput SHAPE-MaP approach for in-cell RNA structural probing and compared structural changes impacted by the CASM34 (c.102 C>A) following expression in the p53 null cell H1299 cells. Compared to p53-WT, the CASM34 mutant significantly altered the secondary structure of the *p53* mRNA, as indicated in circular plots and SuperFold derived secondary structure models **(Figures 1B-E and Supplementary Figure 1)**. In particular, the regions between +170 to +370 of the *p53* mRNA were significantly altered in the base-pairing probabilities. Delta SHAPE (ΔSHAPE) analysis was used to assess differences in the structural constraints between *WT* and *CASM34* RNAs based on the SHAPE reactivity. ΔSHAPE showed that the CASM34 imposed structural constraints including the 5’ part of the coding sequence harboring riboswitch-like structures and protein binding platforms that are critical for DNA-damage induced ATM kinase-mediated p53 activation **(Figure 1F)**. Interestingly, analysis of the second CASM34 (c.102 C>G) mutant derived from malignant melanoma showed a similar secondary structure alteration in the *p53* mRNA as the (c.102 C>A) **(Figure 1G and Supplementary Figure 1)**. Importantly, the non-cancer-associated synonymous variant (c.102 C>T), caused *p53* mRNA structural alterations distinct from the two CASM34 mutants **(Supplementary Figure 1)**.

**Figure 1.**
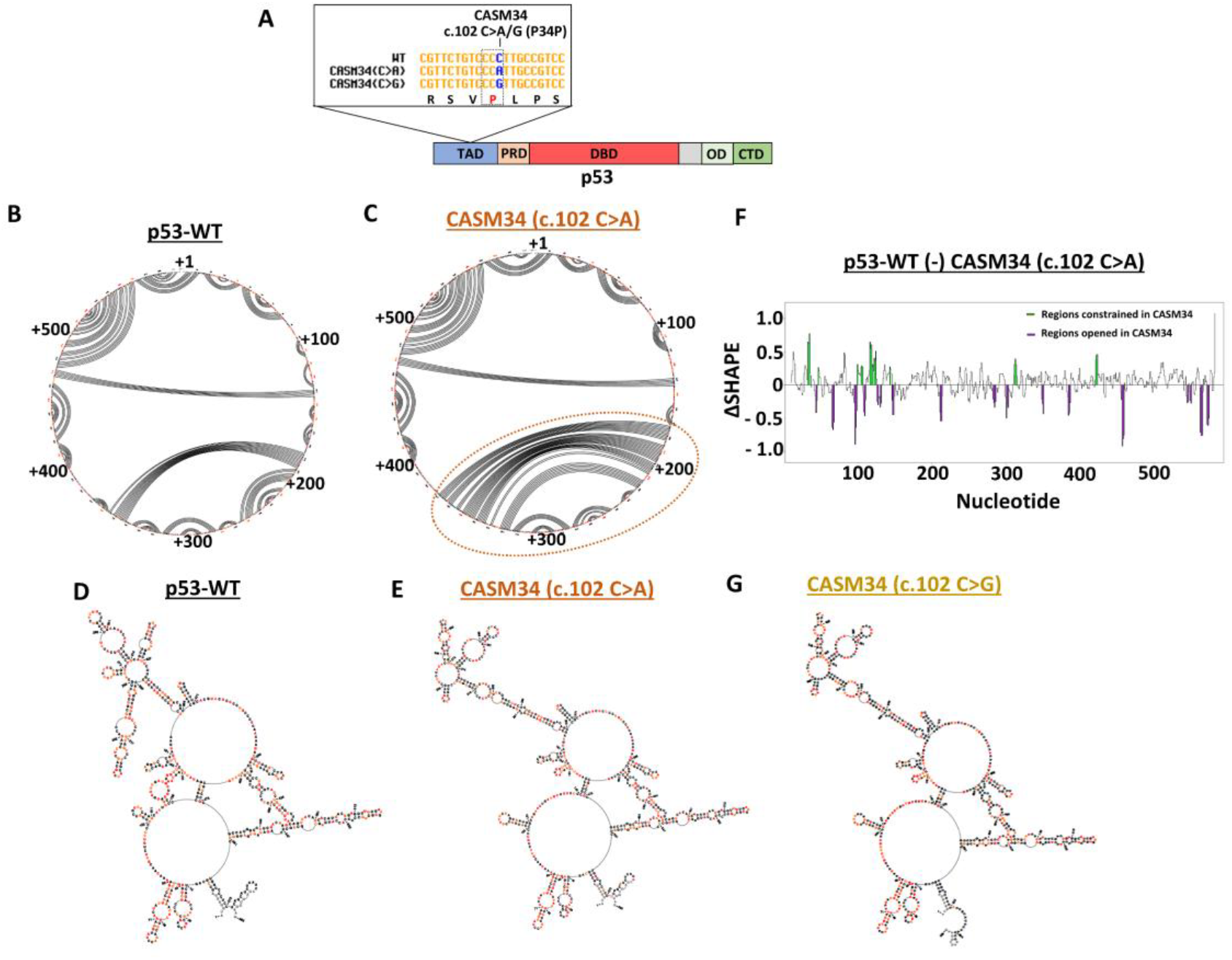
CASM34 alters *p53* mRNA structure in vivo. **A)** Scheme showing the functional domains of p53, with the location of CASM34 mutants indicated in the zoomed-in panel. Transactivation (TAD) domain; proline-rich domain (PRD); DNA binding domain (DBD); oligomerization domain (OD); C-terminal domain (CTD). **B and C)**. Circular plots showing the base-pairing potentials of the *p53* mRNA coding sequence (CDS) based on SHAPE reactivity with (B) p53-WT and (C) CASM34 (c.102 C>A) expressed in p53 null H1299 cells. Base-pairing across the nucleotides are indicated by lines and modifications in the lining pattern indicate RNA secondary structure alterations. RNA Regions which are significantly modified in secondary structure in CASM34 are indicated with dashed orange oval. **D)** and **E)** Secondary structures of indicated *p53* mRNAs coding sequences using the SuperFold algorithm based on SHAPE values. SHAPE modified nucleotide sequences are indicated in orange and red. **F)** ΔSHAPE analysis demonstrates significant structural variations in the *p53* mRNA CDS between p53-WT and CASM34 (c.102 C>A) mutant, with RNA regions constrained in CASM34 (c.102 C>A) are highlighted in green and the regions opened or exposed highlighted in violet. **G)** Same as (D) and (E) based on SHAPE values with CASM34 (c.102 C>G) mutant. SHAPE-Map data shown are representative of at least two independent repeats. RNA secondary structure models of p53-WT, showed in B) and D)are extracted from (15)

### CASM34 prevents DNA damage-induced structural changes in the *p53* mRNA

The two CASM34 mutants cause similar changes in secondary structures of the *p53* mRNA under normal conditions and we next tested if CASM34s affect *p53* mRNA secondary structure in response to genotoxic stress. DNA damage was induced by administering 1 µM of doxorubicin (DOXO) for 8h in H1299 cells expressing p53-WT or CASM34(C>A). In-cell RNA structure probing revealed that the *p53-*WT mRNA successfully induced RNA structural switch under DNA-damage conditions (DOXO). We observed significant alterations in the base pairing pattern of the *p53* mRNA CDS between the RNA regions +1-8 to +105-111 nts., +11-14 to +94-97 nts. and long-range base pairing between regions +110-135 to +575-590 nts. **(Figures 2A and 2B)** (15). Surprisingly, the CASM34 *p53* mRNA showed minimal structural alterations during DNA damage conditions **(Figures 2C-D and Supplementary Figures 2 and 3)**. Thus, a single synonymous mutation in codon 34 prevents the DNA damage response (DDR)-induced *p53* mRNA switch.

**Figure 2.**
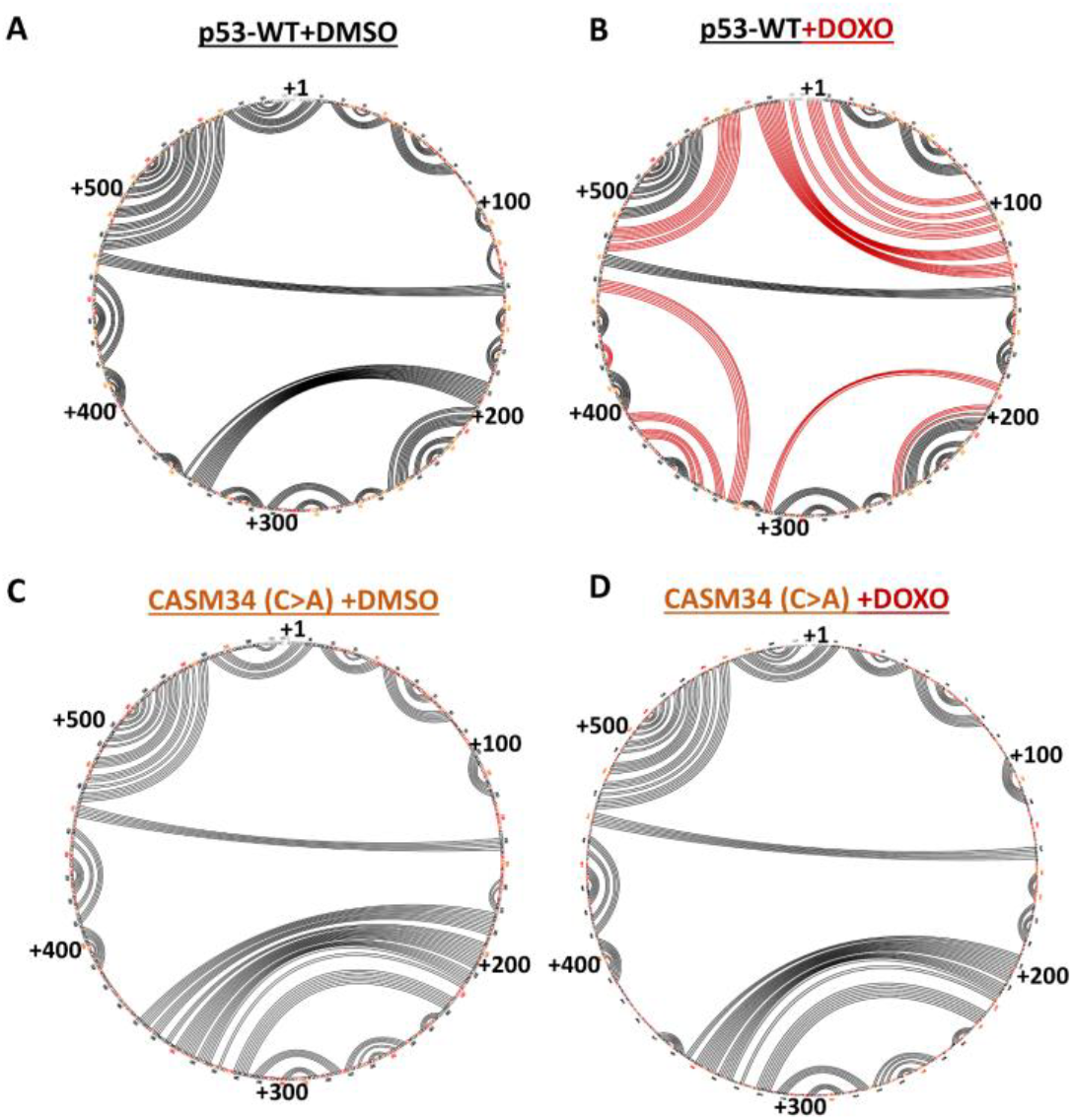
CASM34 targets DNA damage induced *p53* mRNA structural switch. Circular plots showing the secondary structure of *p53*-WT mRNA based on the SHAPE reactivity under **(A)** normal and **(B)** DNA damage conditions (H1299 cells treated with 1 µM doxorubicin (DOXO) for 8h). DNA damage-induced alterations in the mRNA base-pairing patterns are indicated with red lines in the circular plot. **C)** and **D)** circular plots show the secondary structures of *CASM34* mRNA under normal and DNA damage conditions (DOXO). SHAPE modified nucleotide sequences are indicated in orange and red. SHAPE-Map data shown are representative of at least two independent repeats. Circular plots A) and B) are extracted from (15).

### CASM34s affect the cellular DNA damage response pathway

We next sought to determine if CASM34s has physiological impact for the p53 response. We determined the expression profiles of various transcripts in cells expressing CASM34 (c.102 C>A) compared to p53-WT from independent experiments expressing similar levels of p53 protein from either construct **(Figure 3A)**. RT-qPCR was used to determine the expression of a limited set of well-known p53 target genes for cell cycle regulation (*p21*^*CDKN1A*^ *& 14-3-3-σ*) and for apoptosis (*NOXA & PUMA*). Despite expression at similar levels from each mRNA, p53 expressed from the *CASM34* showed a significant lower induction of *PUMA* and *14-3-3-σ* as compared to *p53-WT*, while *NOXA* and the cell cycle arrest factor *p21*^*CDKN1A*^ showed a similar level **(Figure 3B)**. Following 1 µM doxorubicin treatment for 8h, we observed that all four p53 target genes showed an increase in expression, as expected, but the induction of *PUMA* and *14-3-3-σ* were less in cells expressing p53 from *CASM34* **(Figure 3B)**. We also performed ChIP using the p53 antibody (DO-1) under similar conditions followed by qPCR. In agreement with the mRNA expression profiles, we observed that p53 expressed from the *CASM34* mRNA showed reduced binding towards the *PUMA* and *14-3-3-σ* promoters under normal conditions, compared to *p53-WT* mRNA. There was no significant difference in the binding to the *p21*^*CDKN1A*^ promoter **(Figure 3C)**. Similar to CASM34 (c.102 C>A), CASM34 (c.102 C>G) mutant also showed lesser induction on *14-3-3-σ* and *PUMA* expression under normal and DNA damage conditions **(Supplementary Figure 4)**.

**Figure 3.**
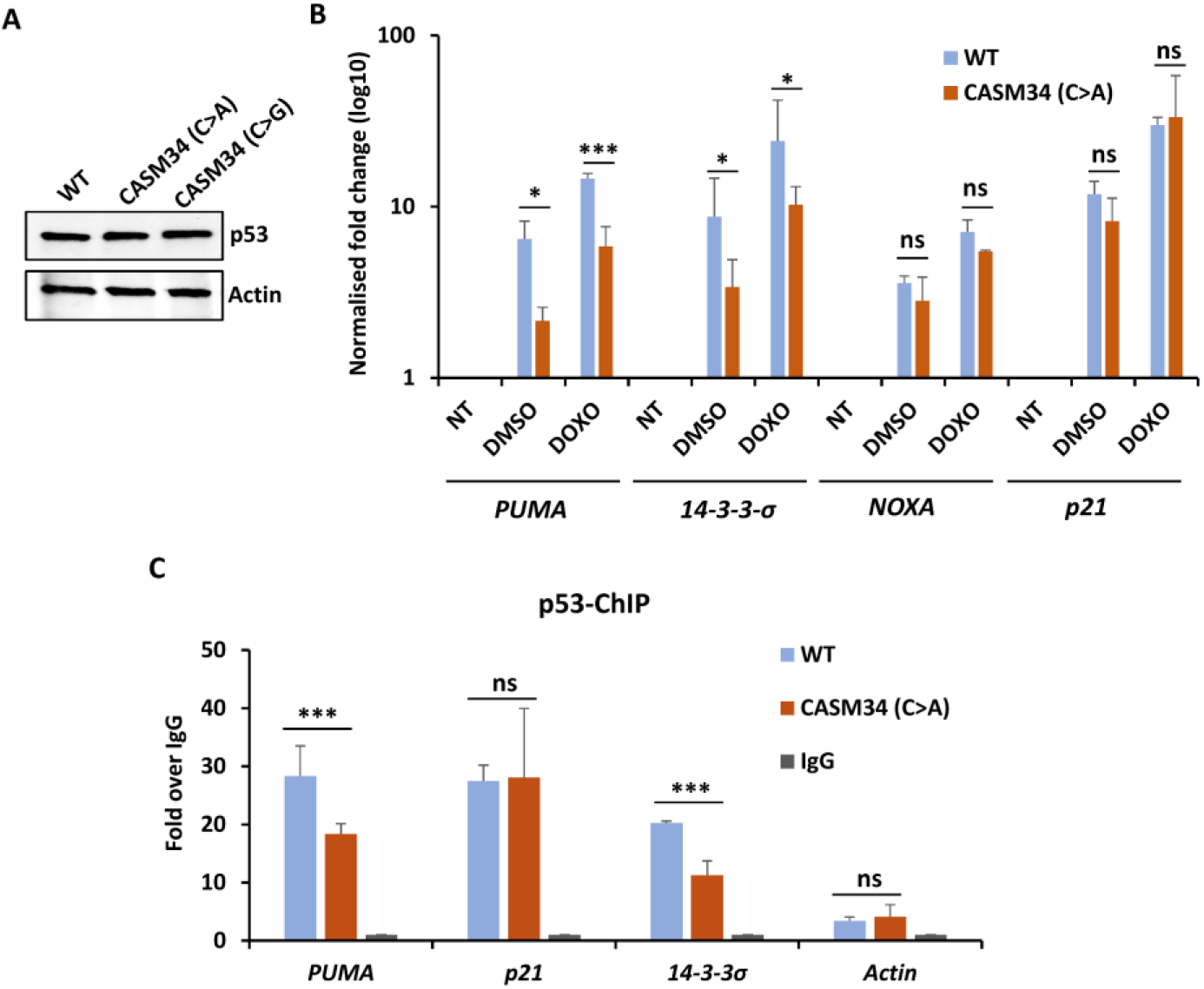
CASM34 selectively affects p53 downstream targets. **A)** Western blot showing p53 levels expressed from indicated cDNA constructs in H1299 cells. Actin was used as a loading control. **B)** Bar graph showing the changes in transcript levels of indicated p53 downstream targets following expression of indicated p53 constructs under normal (DMSO) and DNA damage conditions (1 µM doxorubicin (DOXO) for 8h). **C)** ChIP-qPCR data showing p53 expressed from the WT or CASM34 (c102. C>A) mRNAs binding to indicated downstream target promoters. Relative ChIP signal was normalized with IgG antibody controls, and the corresponding fold enrichments are plotted. For RT-qPCR and ChIP-qPCR, mean ± s.d of three independent experiments are plotted. Statistical significance was calculated using unpaired t-test (***p < 0.001; **p < 0.01; *p < 0.05; ns-not significant).

Despite encoding identical proteins at similar levels, the above data indicate that p53 expressed from the *CASM34 (c*.*102 C>A)* and from the *p53-WT* mRNAs show qualitative differences. We next assessed the effect of CASM34 (c.102 C>A) on genome-wide transcripts by isolating total RNAs from cells expressing WT and CASM34 followed by RNA-sequencing. Gene set enrichment analysis showed altered transcript levels linked to the p53 DNA damage response (DDR), including DNA damage checkpoint signaling and response to ionizing radiation **(Figures 4A-C)**. In parallel, p53 ChIP-seq analysis did not reveal an overall genome-wide change in promoter occupancy between cells expressing p53 proteins from the *WT* or *CASM34* mRNAs **(Supplementary Figures 5A and 5B)**. However, analysis of individual promoters for p53 binding indicated that CASM34 influences p53 promoter occupancy of genes (NOXA, FOXD1, BCL2, PERP, BTF3) which are involved in DDR, p53-mediated apoptosis, and DNA repair processes **(Supplementary Figures 5C-G)**. Importantly, the altered p53-response elements (p53-RE) observed in the ChIP-seq data are in line with the altered expression of the corresponding transcripts observed in genome-wide transcriptome analysis.

**Figure 4.**
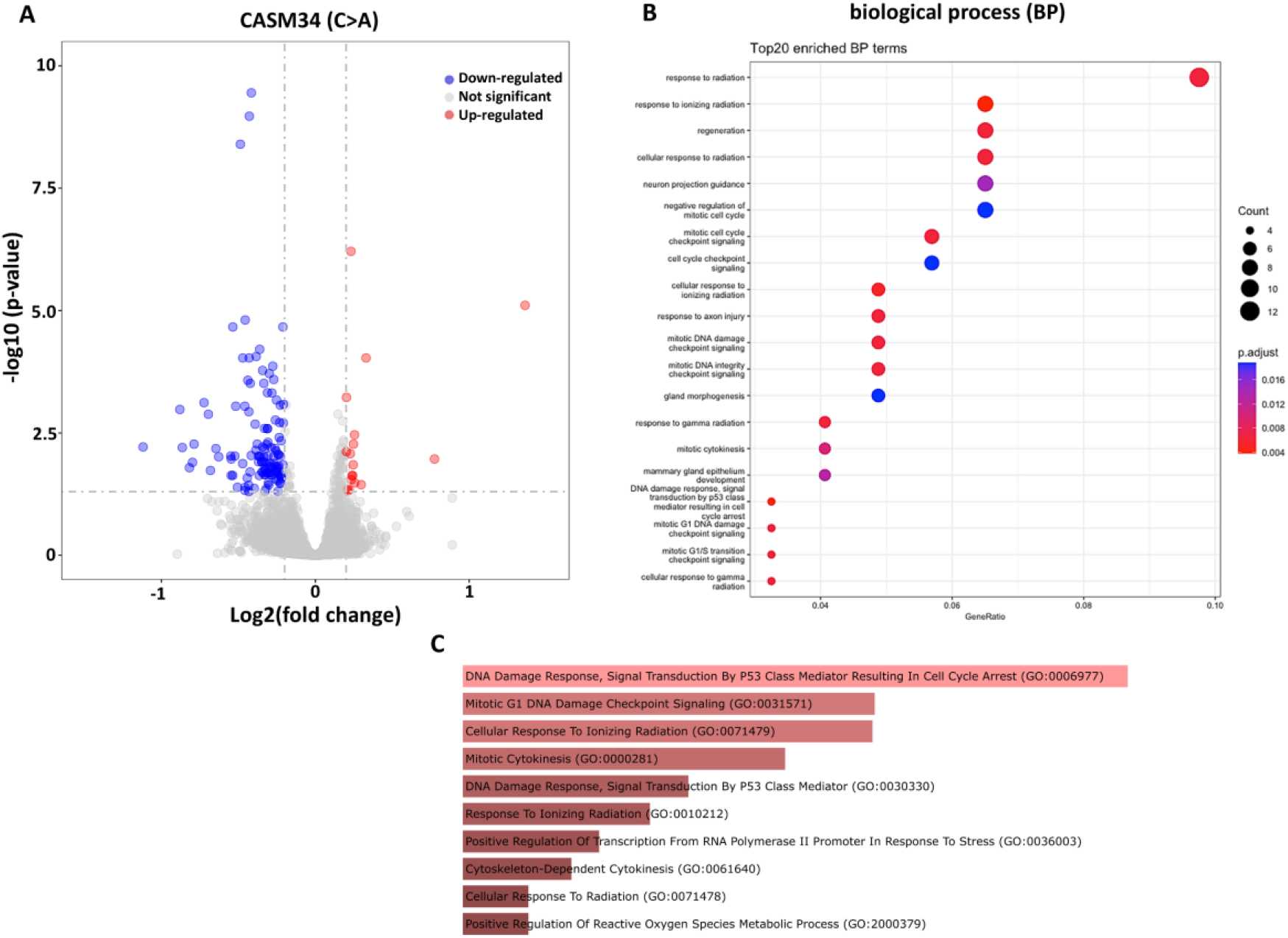
CASM34 modulates transcripts involved in DNA damage response. **A)** Volcano plot showing changes in expression levels of genome-wide transcripts in H1299 cells expressing p53-WT and CASM34 (c. 102 C>A) based on RNA-seq data of at least two independent replicates. **B and C)** Gene enrichment analysis of RNA-sequencing data categorizing the altered transcripts level following expression of p53-WT or CASM34 (c. 102 C>A) mRNAs.

## Discussion

Synonymous mutations (SMs) are emerging as a contributing factor to various diseases, including cancers (9–11, 29). Despite this, SMs are often ignored in genetic analysis and excluded from cancer diagnosis and prognosis. With new information on how SMs affect cell biology and the development of diseases it is expected this will change and that SMs will contribute to personalized medicine (30). We have used the *p53* mRNA as a model to study how RNA structures play an important role in signaling pathways and how SMs can affect the encoded protein and impact on cell biology by interfering with mRNA folding. The two CASMs in codon proline 34 (c.102 C>A and C>G) were identified from malignant glioma and melanoma patients. Both mutants impose distinct, but similar, effects on *p53* mRNA structures, both locally and globally. Remarkably, the two CASM34 single nucleotide changes completely abolish the RNA structural switch observed following DNA damage. That single nucleotide substitutions can abolish the formation of signal-induced mRNA structures is surprising and highlights the highly dynamic nature of *p53* mRNA structures. This is supported by the observation that the *p53* mRNA shows riboswitch-like features during DNA damage and it is plausible that CASM34s interfere with this process **(Figure 2)**(15). Proline is encoded by four codons, and it is interesting to note that the introduction of the non-cancer-associated SM in codon 34 resulted in an RNA structure distinct from the two CASM34s, indicating a selection for either of the two CASM34s during cancer development.

The capacity of the *p53* mRNA to take different conformations in response to different signaling pathways is demonstrated by the observation that a single CASM in codon 203 promotes an *p53* mRNA structure similar to that induced by the PERK kinase during the unfolded protein response, resulting in the expression of the p47 isoform (16),(18). The selective effect of CASMs are illustrated by the fact that PERK-mediated changes in *p53* mRNA structure is not affected by the CASM22, which instead disrupts the formation of an MDM2-binding RNA platform and p53 induction during the DDR (15).

The CASM34 mutants do not affect the steady state p53 protein levels, which is illustrated by the fact that it does not affect the induction of *p21*^*CDKN1A*^. However, global genome-wide analysis showed that that CASM34 (c.102 C>A) primarily affects the expression of genes involved in the p53 downstream DNA damage-response. Genome-wide ChIP-seq analysis detected only a fraction of p53 response elements showing altered p53 binding when synthesized from *p53-WT* or *CASM34* mRNAs. However, ChIP along with sensitive RT-qPCR approach detected selective differences in p53 binding towards response elements when expressed from the CASM34 that agree with the transcript levels of *p21*^*CDKN1A*^, *NOXA, PUMA* and *14-3-3-σ*.

The underlying molecular mechanisms of how mRNA structures contribute to regulating p53 activity are not clear. Two conformation sensitive anti-p53 antibodies (Abs 1620 and 240) did not show any obvious conformational changes between p53 proteins expressed from the different mRNAs **(Supplementary Figure 6)**. It cannot be ruled out that other conformational states might be attained by CASM34 mutants that are not detected by these antibodies. Nevertheless, it is unlikely that the mRNA affects the activity of the encoded protein once the protein has left the ribosome. In line with this notion, previous works have shown that the CASM22 prevents ATM kinase-induced phosphorylation of the nascent p53 protein at serine 15 (13). This is an important step in p53 activation and indicates that mRNA structures play a role in bringing factors to the ribosome that control the activity of the encoded protein. It is, thus, plausible that the difference in activity of p53 synthesized from the *CASM34* mRNA is likewise due to modifications of the nascent p53 protein **(Supplementary Figure 7)**. In this scenario, the CASMs have an indirect effect on the protein, as compared to missense or nonsense mutations that affect the protein directly. We are currently looking into this possibility and, if this turns out to be correct, it makes overexpression a less suitable system for in depth physiological implications of CASMs and further studies using animal models will allow deeper analysis on the physiological effects of CASM34 and its impact on p53 modifications under normal or stress conditions.

## Consent for publication

Not applicable

## Availability of data and materials

The authors declare that data supporting the findings of this study are available within the article and/or in the supplementary information. Raw data of RNA-seq and SHAPE-MaP are available at https://github.com/medbioumu/suppl_casm34

## Competing interests

The authors declare no conflicts of interest.

## Funding and acknowledgments

This work was supported by Cancerfonden, Cancerforskningsfonden Norr, Veteskapsradet, and the Czech Science Foundation, project no. 23-06884S and by MH CZ - DRO (MMCI, 00209805). SVG was supported with grants Cancerforskningsfonden Norr (LP 24-2375; AMP 22-1076), and Basic Science-Oriented Biotechnology Research grant at the Faculty of Medicine, Umeå University (984461). OB was supported with the Ministry of Health of the Czech Republic, grant nr. NW24-10-00204.

## Notes

### Competing Interest Statement

The authors have declared no competing interest.

## References

1. Icgc Tcga Pan-Cancer Analysis of Whole Genomes Consortium, Pan-cancer analysis of whole genomes. Nature 578, 82–93 (2020).

2. I. Cortes-Ciriano, D. C. Gulhan, J. J. Lee, G. E. M. Melloni, P. J. Park, Computational analysis of cancer genome sequencing data. Nat Rev Genet 23, 298–314 (2022).

3. E. R. Kastenhuber, S. W. Lowe, Putting p53 in Context. Cell 170, 1062–1078 (2017).

4. G. Blandino, et al., Mutant p53 protein, master regulator of human malignancies: a report on the Fifth Mutant p53 Workshop. Cell Death Differ 19, 180–3 (2012).

5. A. J. Levine, p53: 800 million years of evolution and 40 years of discovery. Nat Rev Cancer 20, 471–480 (2020).

6. S. Vadivel Gnanasundram, O. Bonczek, L. Wang, S. Chen, R. Fahraeus, p53 mRNA Metabolism Links with the DNA Damage Response. Genes (Basel) 12 (2021).

7. J. V. Chamary, L. D. Hurst, Evidence for selection on synonymous mutations affecting stability of mRNA secondary structure in mammals. Genome Biol 6, R75 (2005).

8. R. C. Hunt, V. L. Simhadri, M. Iandoli, Z. E. Sauna, C. Kimchi-Sarfaty, Exposing synonymous mutations. Trends Genet 30, 308–21 (2014).

9. Y. Sharma, et al., A pan-cancer analysis of synonymous mutations. Nat Commun 10, 2569 (2019).

10. F. Supek, B. Minana, J. Valcarcel, T. Gabaldon, B. Lehner, Synonymous mutations frequently act as driver mutations in human cancers. Cell 156, 1324–1335 (2014).

11. E. Herreros, X. Janssens, D. Pepe, K. D. Keersmaecker, “SNPs Ability to Influence Disease Risk: Breaking the Silence on Synonymous Mutations in Cancer” in Single Nucleotide Polymorphisms: Human Variation and a Coming Revolution in Biology and Medicine, Z. E. Sauna, C. Kimchi-Sarfaty, Eds. (Springer International Publishing, 2022), pp. 77–96.

12. M. M. Candeias, et al., P53 mRNA controls p53 activity by managing Mdm2 functions. Nat Cell Biol 10, 1098–105 (2008).

13. K. Karakostis, et al., A single synonymous mutation determines the phosphorylation and stability of the nascent protein. J Mol Cell Biol 11, 187–199 (2019).

14. L. Malbert-Colas, et al., HDMX folds the nascent p53 mRNA following activation by the ATM kinase. Mol Cell 54, 500–11 (2014).

15. S. Chen, et al., The p53 mRNA exhibits riboswitch-like features under DNA damage conditions. iScience 28, 113555 (2025).

16. L. Fusee, et al., The p53 endoplasmic reticulum stress-response pathway evolved in humans but not in mice via PERK-regulated p53 mRNA structures. Cell Death Differ (2023). 10.1038/s41418-023-01127-y.

17. L. Haronikova, et al., The p53 mRNA: an integral part of the cellular stress response. Nucleic Acids Res 47, 3257–3271 (2019).

18. R. Sajwan, et al., A cancer-associated TP53 synonymous mutation induces synthesis of the p53 isoform p53/47. British Journal of Cancer (2025). 10.1038/s41416-025-03127-w.

19. R. R. Breaker, Riboswitches and the RNA world. Cold Spring Harb Perspect Biol 4 (2012).

20. R. P. Smyth, M. Negroni, A. M. Lever, J. Mak, J. C. Kenyon, RNA Structure-A Neglected Puppet Master for the Evolution of Virus and Host Immunity. Front Immunol 9, 2097 (2018).

21. C. J. Lewis, T. Pan, A. Kalsotra, RNA modifications and structures cooperate to guide RNA-protein interactions. Nat Rev Mol Cell Biol 18, 202–210 (2017).

22. K. M. Weeks, SHAPE Directed Discovery of New Functions in Large RNAs. Acc Chem Res 54, 2502–2517 (2021).

23. M. J. Smola, G. M. Rice, S. Busan, N. A. Siegfried, K. M. Weeks, Selective 2’-hydroxyl acylation analyzed by primer extension and mutational profiling (SHAPE-MaP) for direct, versatile and accurate RNA structure analysis. Nat Protoc 10, 1643–1669 (2015).

24. M. J. Smola, K. M. Weeks, In-cell RNA structure probing with SHAPE-MaP. Nat Protoc 13, 1181–1195 (2018).

25. T. Wu, et al., clusterProfiler 4.0: A universal enrichment tool for interpreting omics data. Innovation (Camb) 2, 100141 (2021).

26. M. V. Kuleshov, et al., Enrichr: a comprehensive gene set enrichment analysis web server 2016 update. Nucleic Acids Res 44, W90–7 (2016).

27. B. Leroy, et al., The TP53 website: an integrative resource centre for the TP53 mutation database and TP53 mutant analysis. Nucleic Acids Res 41, D962–9 (2013).

28. M. Miladi, M. Raden, S. Diederichs, R. Backofen, MutaRNA: analysis and visualization of mutation-induced changes in RNA structure. Nucleic Acids Res 48, W287–W291 (2020).

29. Z. E. Sauna, C. Kimchi-Sarfaty, Understanding the contribution of synonymous mutations to human disease. Nat Rev Genet 12, 683–691 (2011).

30. Y. Lan, et al., Synonymous mutations promote tumorigenesis by disrupting m6A-dependent mRNA metabolism. Cell 188, 1828-1841.e15 (2025).

